# Alzheimer’s Disease Brain Phenotypes are Age-dependent

**DOI:** 10.64898/2026.03.31.715296

**Authors:** Fermin Travi, Anushree Mehta, Eduardo Castro, Hongyang Li, Jenna Reinen, Amit Dhurandhar, Pablo Meyer, Diego Fernández Slezak, Guillermo A. Cecchi, Pablo Polosecki

## Abstract

A widespread view of neurodegenerative disorders, including Alzheimer’s Disease (AD), frames their effects as accelerated aging, with the brain-age gap (BAG, the deviation of predicted ‘brain age’ from chronological age) as a staple biomarker. However, BAG relies on a fundamental, untested assumption: that AD can be identified via age-invariant brain phenotypes. Using invariant representation learning on brain MRI from 44,178 individuals, we created neural representations that optimally convey age information (age-aware) or conversely remove it (age-invariant) while minimizing reconstruction distortion. We provide the first causal evidence that age information is necessary in brain biomarkers for AD detection: age-aware representations achieve competitive state-of-the-art performance and significantly outperform age-invariant ones (0.84 vs. 0.77 AUC, p < 0.001, with external validation). This necessity reveals a conceptual flaw in BAG: by subtracting chronological age, it discards the very information essential for accurate detection. Using conditional decoders to simulate aging trajectories, we found that healthy aging and AD operate along multiple independent anatomical dimensions (deep gray matter, frontoparietal, temporal). AD patients diverge from rather than accelerate healthy aging, showing pathological temporal shifts alongside, remarkably, relative frontoparietal preservation. Furthermore, representational similarity analysis suggests that even models pretrained on non-age tasks (e.g., sex or BMI) implicitly converge toward age-related features when optimized for AD. Given that the AD phenotype cannot be decoupled from age, our results establish a hard limit for age-independent biomarkers and favor multidimensional models that preserve aging structure over unidimensional summaries like BAG.

## 1. Introduction

Neurodegeneration involves progressive structural and functional deterioration of neurons culminating in cell death, encompassing a range of biological processes^1,2^. Whether neurodegenerative disease represents a qualitatively distinct pathological process or an exacerbation of normal aging mechanisms remains unresolved, with evidence supporting both a continuum model and categorical distinctions based on disease-specific molecular drivers and selective neuronal vulnerability patterns^3–5^. The robust predictability of chronological age from brain images^6,7^ has elevated the conceptualization of neurodegeneration as a deviation from healthy aging in the imaging field. This framing has been central to recent efforts to extract diagnostic and prognostic biomarkers using machine learning techniques^8^, driving attempts to detect neurodegeneration, particularly in Alzheimer’s Disease (AD)^9^. These efforts are based on the hypothesis that the effects of the disorders being studied manifest in images as a form of premature or accelerated aging^10^, and it constitutes the base of notable machine learning applications. Given its history, it is remarkable that basic assumptions behind this hypothesis and its applications have not been tested against suitable alternatives.

The brain aging hypothesis was formulated around the notion that the difference between estimated brain age and chronological age (the brain-age gap, BAG) is a biomarker for degeneration and other disorders^9,11,12^. BAG is meant to quantify the degree of deviation from normative aging in an individual (Fig. 1 A). It comes with limitations. First, it is directly based on the error of a predictive model, encapsulating its biases and effects of non-aging-related heterogeneity and measurement noise^13^. Moreover, BAG is a one-dimensional number, whereas aging is a high-dimensional process^14,15^ (Fig. 1 B). It has been proposed, therefore, that the use of latent features from age prediction models, with or without fine-tuning^16,17^, more accurately captures the effects of disease by preserving aging information. However, an alternative view argues that more effective group identification emerges from disentangling target phenotypes from their degree of progression^18,19^. BAG itself embodies this latter approach: by subtracting chronological age from estimated brain age, it constructs a measure that is by design age-invariant, with direct attempts to further maximize this property^20^. These conflicting approaches raise a fundamental, untested question: is the aging signal itself necessary for detecting neurodegeneration, or should it be removed to reveal static, disease-specific footprints?

**Figure 1:**
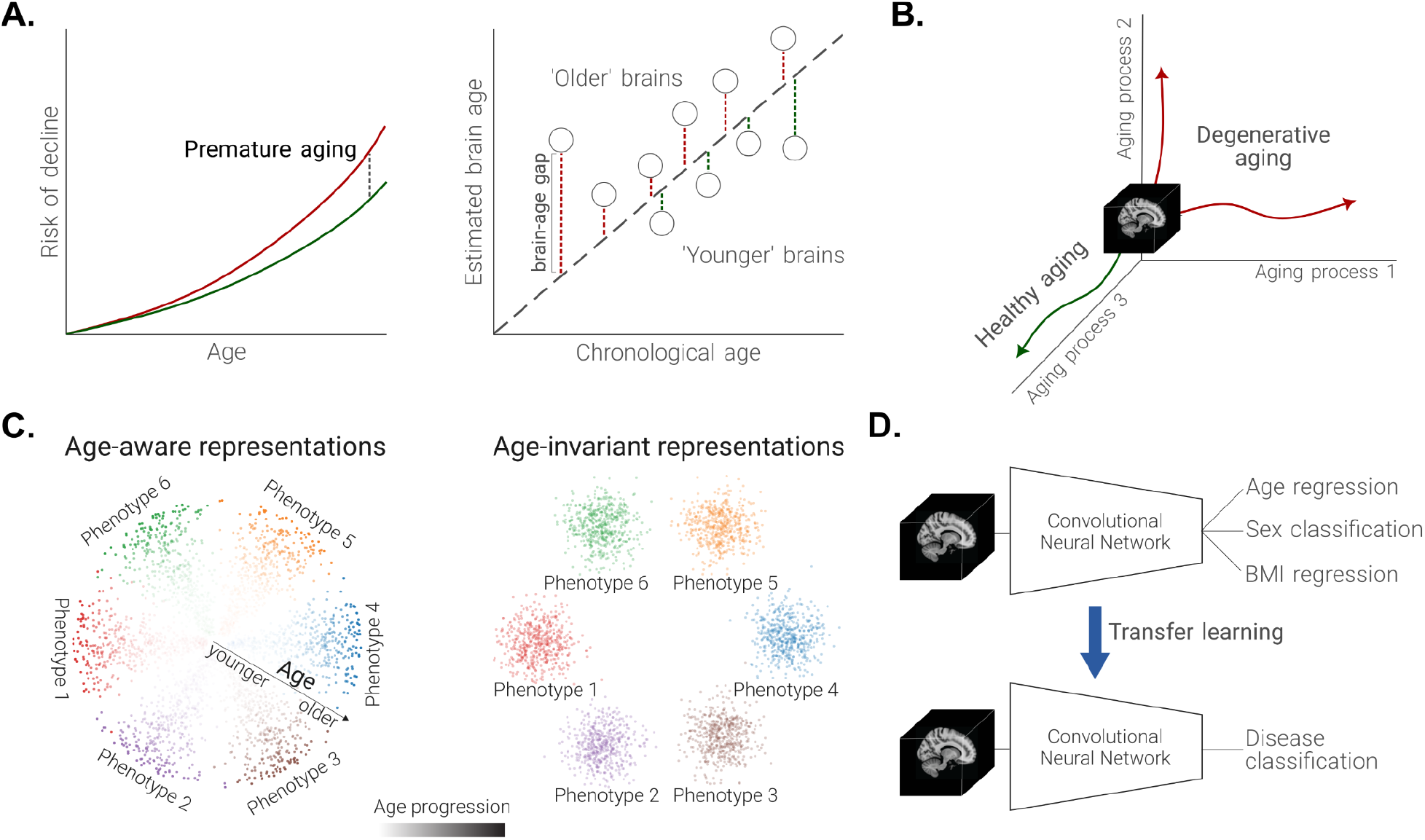
**A**. Unidimensional account of the brain aging hypothesis. On the left, cumulative tissue damage over time increases disease risk, making individuals with above-average damage effectively age prematurely. On the right, the brain-age gap (BAG), defined as the age estimation error of a predictive model, is thought to reflect this excess accumulation. **B**. Multidimensional view of brain aging. Rather than a single timeline, aging is conceptualized as a high-dimensional manifold shaped by multiple independent biological processes. Under this framework, neurodegenerative disease does not merely accelerate normative aging, but drives the brain along distinct, pathological trajectories that actively diverge from healthy structural patterns. **C**. Conceptualization of latent brain representations with and without age information. Each dot represents a subject. In the age-aware representation (left), different phenotypes become more distinct as they evolve, and brain age can be read out. Age-invariant representations (right), conversely, remove progression information, potentially isolating static disease phenotypes. **D**. Representational shifts in transfer learning. To isolate the specific value of age information, models are pre-trained on diverse phenotypic tasks (age, sex, and BMI) and subsequently fine-tuned for disease classification. By measuring the representational distance before and after fine-tuning, we evaluate how much each network’s internal structure must shift to detect neurodegeneration, testing whether age-pretraining inherently aligns with pathological signatures.

Deep learning approaches offer unprecedented opportunities to answer this question by building models that systematically manipulate the information represented, enabling researchers to add or remove specific types of information^21–23^. This allows causal testing of aging information’s contribution to predictive inference. Specifically, one can construct representations that are age-agnostic (naive unsupervised), age-invariant (deliberately removing aging information while preserving other morphological and phenotypic information), or age-aware (maximally preserving aging information). Age-aware representations align with the hypothesis that aging information enables disease detection, while age-invariant representations align with the BAG framework by learning brain aspects that remain stable over time, representing aging styles rather than aging degree (Fig. 1 C). A first contribution of this study, then, is to directly test whether aging information is necessary for detecting neurodegeneration by learning age-agnostic, age-invariant, and age-aware representations from large population datasets and testing their application to dementia detection in case-control cohorts.

One major recent thrust of the brain aging hypothesis uses transfer learning to move beyond unidimensional BAG. Networks pretrained on age prediction are fine-tuned for AD detection^16,17,24,25^, with the rationale that aging consists of multiple independent processes^14,15^ that intermediate representations express more richly than BAG^17^. However, reported performance improvements of age-pretrained models could simply reflect the benefit of exposure to brain image statistics and its rich phenotypic information, regardless of pretraining task^26^. The literature is equivocal: some studies show advantages for age-based pretraining^16^, others employed unrelated pretraining successfully without considering age prediction^26^, and still others find no-pretraining performs equally well^27^. This creates a paradox: why is high performance possible in such disparate conditions? Our second contribution is to study the effect of brain-based pretraining tasks (age, sex, BMI) and no-pretraining for AD detection transfer learning, resolving this inconsistency by examination of the internal representations learned from the different pretraining tasks and their change during fine-tuning (Fig. 1 D).

Here, we leverage large population-based (n=44,178) and AD-focused case-control studies (n=4,280) to address a fundamental question: is the aging signal inherently necessary for detecting neurodegeneration, or should temporal progression be removed to isolate static disease footprints? Using invariant representation learning tools^28,29^, we demonstrate that age-aware representations outperform age-invariant ones in AD detection, supporting the brain aging hypothesis. However, simulating longitudinal brain change reveals multiple aging dimensions where neurodegeneration distinctly diverges from healthy patterns, adding crucial nuance to this framework. Furthermore, comparing transfer learning approaches (age, sex, and BMI) for disease detection shows that while all pretraining tasks yield similar early-layer representations, age-pretraining uniquely positions networks closer to their fine-tuned goal state. Together, our findings characterize the AD phenotype as inextricably anchored in the aging process. This reveals the fundamental inadequacy of age-independent summaries like BAG, establishing a mandate for multidimensional biomarkers that leverage the essential temporal structure of neurodegeneration.

## 2. Results

We tested the brain aging hypothesis in steps. First, we assembled two composite datasets: a population-based dataset for representation learning and model pretraining, and a case-control dataset for developing and evaluating disease detection models using the learned representations. Second, we designed variational autoencoder (VAE) architectures with three approaches to age information: age-agnostic (no explicit age treatment), age-aware (maximally informed), and age-invariant (maximally uninformed). We evaluated the resulting brain representations on chronological age estimation and brain image reconstruction in the general population, and used them to train and evaluate disease detection models in the case-control dataset. To understand the learned aging patterns, we leveraged the conditional decoder from our age-invariant model to simulate longitudinal aging trajectories and decomposed them into constituent patterns. We then used these to explore aging divergences in case-control groups. Finally, we pretrained models with the same architecture as the encoder portion of the VAEs to estimate age, sex, or BMI, then fine-tuned each for disease detection. We conducted representational similarity analysis (RSA) to quantify representational shifts induced by fine-tuning.

### 2.1 An aggregated dataset for evaluation of aging and disease detection hypotheses

#### General population

Brain images collected from the general population served for model pretraining and learning of abstract brain representations. It comprised 44,482 brain images (mean age = 54.5 years, std. = 8.8; 23,422 females), collected from six studies: UK Biobank (UKBB)^30^, Cam-CAN^31^, Human Connectome Project (HCP)^32^, Aging Human Connectome Project (HCP-Aging)^33^, NKIRS^34^, and SALD^35^. Each image corresponds to a unique participant, using the baseline scan when more than one image was available. The number of participants and their demographic information are summarized in Table 1. Age and sex were available for all cohorts, whereas body mass index (BMI) was only available for UKBB, HCP, and HCP-Aging.

**Table 1:** Cohorts that make up the General population dataset. Each brain image corresponds to a unique individual.

| Dataset | Number of brain images<br>(female ratio) | Age range (mean and SD) |
| --- | --- | --- |
| UK-Biobank | 42,035 (52%, 22,000) | 44.23-82.74 (64.05 ± 7.70) |
| Cam-Can | 455 (52%, 237) | 20.17-87.92 (47.13 ± 14.85) |
| HCP | 661 (56%, 369) | 22.00-36.00 (28.81 ± 3.66) |
| HCP-aging | 403 (52%, 209) | 36.00-89.83 (60.07 ± 16.14) |
| NKIRS | 586 (67%, 393) | 21.00-85.00 (47.46 ± 17.81) |
| SALD | 342 (62%, 214) | 21.00-80.00 (49.36 ± 17.09) |

#### Diseased and healthy controls

The *Diseased* dataset totaled 4115 brain images (mean age 72.3, std. 7.7; 2118 females; 565 AD, 1493 MCI, 2057 Healthy Controls, HC), combined from three studies: ADNI^36^, OASIS^37^, and AIBL^38^. All three have diagnosed each individual as having AD, MCI, or being a HC. The number of brain images and their demographics can be found in Table 2 (see Appx. Fig. A1 for the overall age distribution across diagnosis).

**Table 2:** Cohorts that make up the Diseased dataset. Each brain image corresponds to a unique individual and each of them is diagnosed as one of Alzheimer’s Disease, Mild Cognitive Impairment, or Healthy Control.

| Dataset | Number of brain images (female ratio) | Age range (mean and SD) | Diagnosis |  |  |
| --- | --- | --- | --- | --- | --- |
|  |  |  | AD | MCI | HC |
| ADNI | 2,527 (48%, 1,227) | 50.50-91.50 (72.92 ±7.42) | 421 | 1,167 | 939 |
| OASIS | 905 (56%, 504) | 42.66-93.93 (70.27 ±8.83) | 62 | 193 | 650 |
| AIBL | 683 (57%, 387) | 55.00-93.00 (72.91 ±6.55) | 82 | 133 | 468 |

### 2.2 Aging information is necessary for neurodegeneration detection

To causally test the brain aging hypothesis we implemented three different VAE architectures: *age-*agnostic, which worked as a standard VAE; *age-*aware, designed to maximally embed age information in the encodings; and *age-invariant*, designed to explicitly remove age information from the encodings (see Methods) (Fig. 2 A). We additionally developed architectures that either incorporated or excluded other phenotypic data (such as sex or BMI) to serve as control models (Appendix A2).

**Figure 2:**
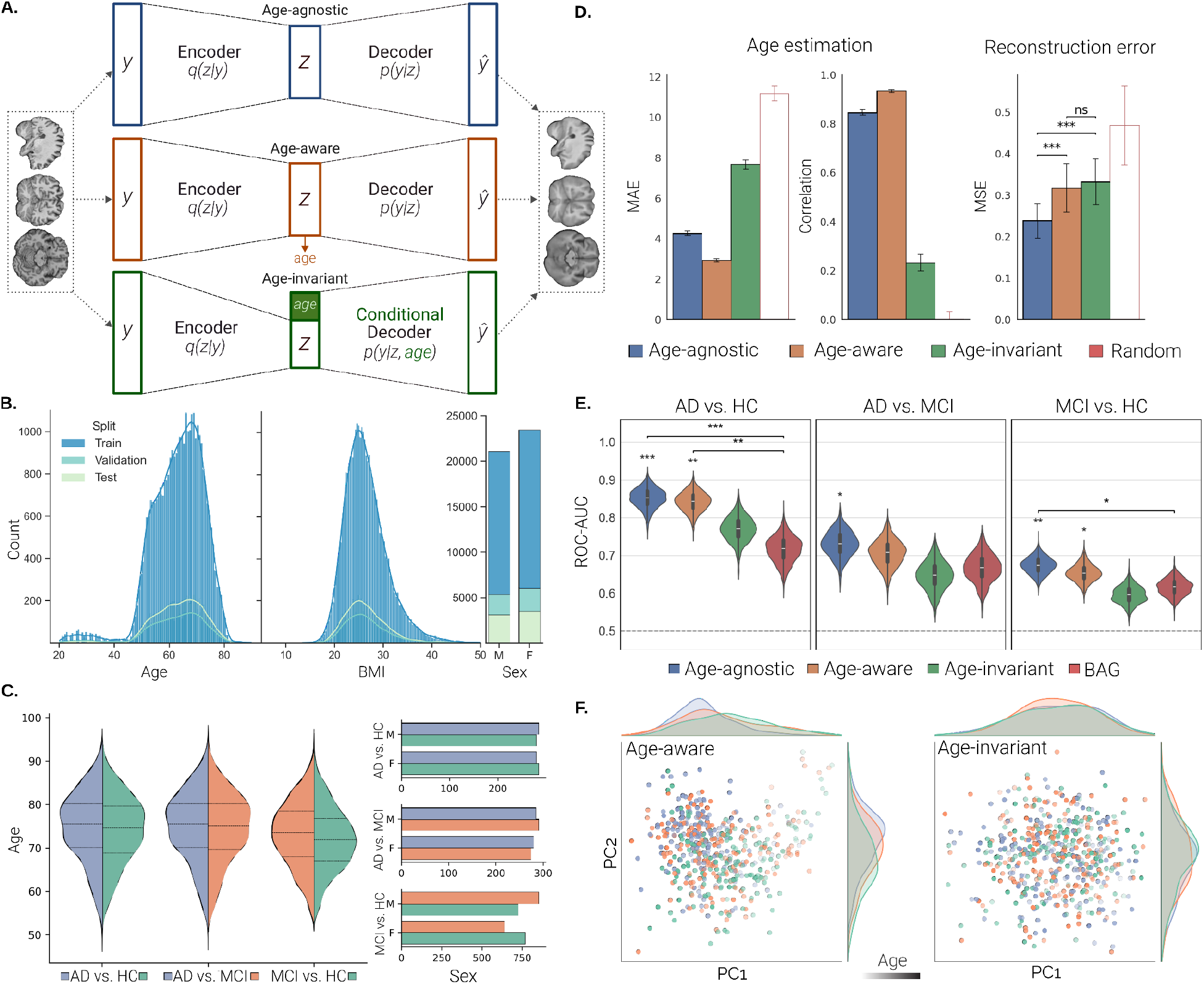
**A**. Schematic of the variational autoencoders, from top to bottom: age-agnostic, age-aware, and age-invariant. **B**. Age, BMI and sex distributions across splits in the General population dataset. **C**. Age and sex distribution in each of the three disease classification tasks (Alzheimer’s Disease vs. Healthy Controls, Alzheimer’s Disease vs. Mild Cognitive Impairment, Mild Cognitive Impairment vs. Healthy Controls) after matching by age and sex. **D**. Age estimation and reconstruction error from the different brain representations (age-invariant, age-agnostic, age-aware) on the test set of the General population dataset. Age estimation error is quantified using Mean Absolute Error (MAE) and Pearson’s correlation. The random baseline is constructed by shuffling the labels. The reconstruction error corresponds to the Mean Squared Error (MSE) between the input (T1-w images) and the output of the decoder (reconstructed T1-w images). The values are the mean and the bars the standard deviation of a bootstrapped distribution (N=1,000). Statistical significance between two distributions is determined by the proportion of lower values. **E**. Performance on the detection of Alzheimer’s Disease against Healthy Controls (first from the left), Alzheimer’s Disease and Mild Cognitive Impairment (middle), and Mild Cognitive Impairment and Healthy Controls (third from the left) using age-agnostic, age-aware and age-invariant brain representations and the brain age-gap (BAG), bootstrapped a thousand times. The violin plots reflect the Kernel Density Estimate (KDE) and the median and interquartile range of the ROC-AUCs distribution. Asterisks above the violins indicate statistical significance against age-invariant, determined by the proportion of lower values. **F**. First two principal components of the *age-aware* and *age-invariant* brain representations on a random 30% (N=508) sample of the matched brain images. Marker transparency is determined by the age of the individual.

The population-based dataset was split in 75:10:15 ratios for model training, validation and testing, respectively (Fig. 2 B). To account for the sample size differences between studies, images from smaller studies were upsampled during training. After training, the validation and test sets were encoded to brain representations, and a fully-connected network with 2 hidden layers was fit to them in order to assess the amount of age information they contained (see Appendix A2 for sex and BMI estimation performance as well). Image reconstruction error was measured as the mean squared difference between the input and output of the VAEs. Randomized true labels and the mean brain image served as baselines for age estimation and reconstruction error, respectively. The validation set was used for hyperparameter tuning and the final performance is reported on the test set (see Methods). The resulting best-performing encoding models were then evaluated for disease detection on the Diseased dataset (Fig. 2 C).

Figure 2 D presents the performance of brain representations learned by the aforementioned VAEs in age estimation (measured with MAE and Pearson’s correlation coefficient) and image reconstruction (measured with MSE) on the General population test set. Age estimation is best for *age-aware* (mean MAE = 2.92 years, std = 0.07, r = 0.93), followed by *age-agnostic* (mean MAE = 4.26, std = 0.11, r = 0.85) and *age-invariant* (mean MAE = 7.66, std = 0.21, r = 0.23). All of Pearson’s correlations were significant (p < 0.001). The random baseline mean MAE is 11.16 years (std = 0.37). When it comes to reconstruction performance on the T1-w images, the lowest mean MSE (0.24, std = 0.04) is achieved by *age-agnostic*, followed by *age-aware* (0.32, std = 0.06) and *age-invariant* (0.33, std =0.05). The difference between *age-aware* and *age-invariant*, however, is not significant. *Age-agnostic*, in particular, possesses a lower reconstruction error due to having one less loss term in comparison with the other models (see Methods). The three of them were significantly different with the random baseline, whose mean MSE is 0.47 (std = 0.10). Overall, these results demonstrate that while the three models encode different levels of age-related information, they all retain sufficient structural information to effectively reconstruct brain images.

We then evaluated these encoding models on the Diseased dataset. We considered three classification tasks: AD vs. HC, AD vs. MCI, and HC vs. MCI. Because these groups differed significantly in age (Appendix Fig. A1), we controlled for confounding by matching each individual in the minority class (e.g., AD) with a sample from the majority class (e.g., HC) based on age and sex. This yielded balanced, equally sized groups matched on the variables of interest, at the cost of reducing the number of samples available from the majority group (Table 3 and Fig. 2 C). A logistic regression fit to the brain age-gap (BAG) of individuals is added for comparison (see Methods).

**Table 3:** Number of brain images, mean age and sex for each of the disease classification tasks. AD refers to Alzheimer’s Disease, MCI to Mild Cognitive Impairment and HC to Healthy Controls.

| Task |  | AD | MCI | HC |
| --- | --- | --- | --- | --- |
| AD vs. HC | N | 565 | - | 565 |
| | Mean age (SD) | 74.83 ($\pm 7.95$) | - | 74.13 ($\pm 7.71$) |
|  | Number of females | 280 (49%) | - | 285 (50%) |
| AD vs. MCI | N | 565 | 565 | - |
| | Mean age (SD) | 74.83 ($\pm 7.95$) | 74.51 ($\pm 7.78$) | - |
|  | Number of females | 280 (49%) | 274 (48%) | - |
| HC vs. MCI | N | - | 1,493 | 1,493 |
| | Mean age (SD) | - | 73.20 ($\pm 7.48$) | 71.99 ($\pm 7.07$) |
|  | Number of females | - | 640 (43%) | 768 (51%) |

Among the three classification tasks, the largest performance differences emerged when comparing Healthy Controls (HC) to disease groups (Fig. 2 E). For the AD versus HC distinction (Fig. 2 E, left-most column), *age-agnostic* achieved the highest median AUC of 0.85 (IQR 0.03), followed closely by *age-aware* (median AUC 0.84, IQR 0.03). Both of these significantly outperformed *age-invariant* and BAG, which achieved median AUCs of 0.77 and 0.72 (IQR 0.04), respectively (p < 0.001 and p < 0.01). The AD versus MCI classification (Fig. 2 E, middle column) showed a similar pattern: *age-agnostic* and *age-aware* performed best (0.73 and 0.71 median AUC; IQR 0.05), while BAG and *age-invariant* performed worse (median AUC 0.67 and 0.65; IQR 0.05), with a statistically significant difference between *age-agnostic* and *age-invariant* (*p < 0*.05). Finally, for the HC versus MCI distinction (Fig 2 E, right-most column), *age-agnostic* and *age-aware* again demonstrated superior performance (median AUC 0.67 and 0.65; IQR 0.03), with both significantly outperforming *age-invariant* (median AUC 0.57; IQR 0.02; *p < 0.01* and *p < 0.05*, respectively); *age-agnostic* also significantly outperformed BAG (median AUC 0.62; IQR 0.03; *p < 0.05*). The performance achieved by our age-aware and age-agnostic models on external datasets is effectively on par with recent rigorous benchmarks on these tasks^39^. Consistent with these classification results, principal component analysis of *age-aware* brain representations revealed that the first principal component captures a gradient from HC through MCI to AD (Fig. 2F, left), suggesting these representations organize disease states along a continuum that naturally incorporates aging information. In contrast, age-invariant representations (Fig. 2F, right) show no systematic disease-related ordering—diagnostic groups overlap substantially in the reduced space. This demonstrates that removing age information doesn’t merely reduce signal-to-noise ratio; it disrupts the representational geometry needed to distinguish disease states (see also Appx. Fig. A2 C).

Altogether, these results indicate the relevance of age information when detecting Alzheimer’s Disease and Mild Cognitive Impairment in T1-w brain images. Both *age-agnostic* and *age-aware* performed significantly better when compared to brain representations with reduced age information (*age-invariant* and BAG). Furthermore, in spite of the significant difference in reconstruction error between *age-agnostic* and *age-aware* (Fig. 2 D), these performed comparably in all tasks. This reveals an important principle: the information structure relevant for disease detection is largely orthogonal to general image reconstruction quality. *Age-aware* representations sacrifice some voxel-level reconstruction fidelity to explicitly preserve age-related structural variance, yet this trade-off doesn’t impair disease detection. This suggests that AD-related brain changes are inherently intertwined with age-related patterns, making age information valuable regardless of how explicitly it is modeled.

### 2.3 Multidimensionality in brain aging

To understand what drives dementia detection, we leveraged a key capability of our age-invariant model: its conditional decoder can generate brain images at arbitrary ages while holding other phenotypic factors constant (see Methods). This makes it possible to simulate longitudinal aging trajectories for any individual, effectively asking ‘what would this 45-year-old’s brain look like at 85?’ Using these simulated trajectories, one can decompose the learned aging process into its constituent patterns.

Principal component analysis of the aging differences revealed that the first three PCs explained 10%, 5%, and 4% of the variance, respectively, with a sharp drop thereafter: the first 30 PCs collectively account for only 49% of the total variance in the distribution (Fig. 3A). This fundamentally challenges unidimensional conceptualizations of brain aging. Rather than a single ‘aging vector’ along which all individuals travel at different speeds (the implicit assumption of BAG), these results suggest aging manifests through multiple semi-independent processes that vary in their expression across individuals.

**Figure 3:**
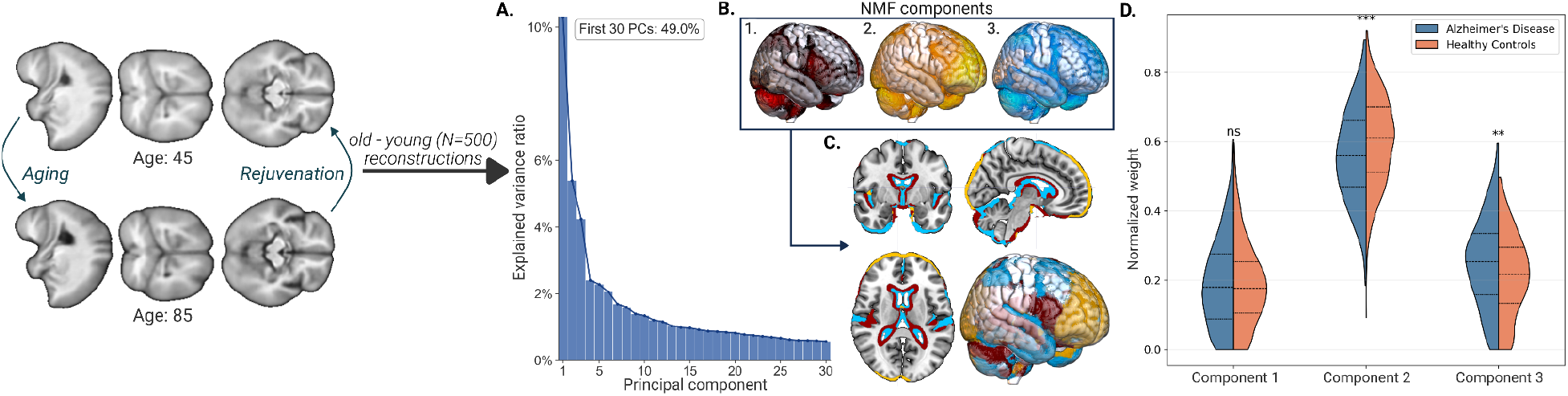
Pipeline to study learned aging information. Brain images corresponding to individuals in the 45-60 and 70-85 age ranges were randomly selected from the test set and divided in groups (young and old, respectively). Their brain scans were reconstructed with the conditional decoder of the *age-invariant* model described in section 2.2. Individuals in the young group had their brain images reconstructed at age 85 and those in the old group at age 45. The relative differences between the older and younger reconstructions were computed and subsequently warped to the MNI template. **A**. PCA components of the relative differences ranked by their percent explained variance. Explained variance decayed sharply after three components. **B**. First three NMF components of the relative differences. **C**. Voxel-wise assignment to the NMF component with the highest value after L2 normalization, thresholded at 10^-3^. **D**. Weight distribution of the NMF components in B. when reconstructing aging differences in Alzheimer’s Disease patients (blue) and Healthy Controls (coral), cut at zero since there are no negative values.

We then explored aging-induced atrophy. Given that the differences between aged and younger reconstructions predominantly reflect unidirectional gray matter loss, non-negative matrix factorization (NMF) and its clustering properties offer the ideal framework to decompose these age-related losses into additive constituent patterns. We extracted three components for this analysis (Fig. 3B; see Methods), as the initial PCA indicated that the first linear three factors captured the most significant proportion of the explained variance. This decomposition revealed three overlapping but distinct patterns of atrophy (Fig. 3C): Component 1 (red) captures deep gray matter loss and ventricular expansion; Component 2 (yellow) reflects frontoparietal and occipital cortical thinning; Component 3 (blue) shows temporal lobe degeneration. Critically, these patterns are not mutually exclusive (Fig. 3C)—most voxels show contributions from multiple components, suggesting they represent partially overlapping biological processes rather than discrete subtypes. The consistency of these patterns with those identified in recent deep learning decompositions^23^ validates that our age-invariant model has learned biologically meaningful aging representations.

Given that these components represent general aging processes, we utilized them to investigate whether these general patterns are differentially expressed in neurodegenerative conditions. We reconstructed brain images from elderly Alzheimer’s Disease (AD) patients and healthy controls (HC) as if they were 45 years old, computed their aging difference maps, and analyzed the contribution of each component (see Methods). As shown in Figure 3 D, a significant difference was observed in the weight distribution of the NMF components when reconstructing these aging differences. The third component, which encompasses temporal lobe atrophy, was significantly more predominant in AD patients (mean 0.25, std. 0.13) than in HC (mean 0.21, std. 0.12; p < 0.01). Conversely, the second component was less predominant in AD patients (mean 0.56, std. 0.13) compared to controls (mean 0.60, std. 0.13; p < 0.001). This observation is consistent with known early neurodegeneration in AD, suggesting that healthy and neurodegenerative aging follow divergent patterns. Collectively, these results demonstrate that brain aging unfolds along multiple dimensions^14,15^, and that the trajectories through this multidimensional space differ between healthy individuals and those with neurodegenerative disease.

### 2.4 Neural networks implicitly learn aging features in neurodegeneration detection

We next investigated an apparent paradox in the literature regarding transfer learning for disease classification: while some studies report superior performance when pretraining on age prediction^16^, others find comparable performance using unrelated pretraining tasks^26^, and still others observe no benefit from pretraining at all^27^. If aging information is truly necessary, why does performance sometimes remain high across such disparate pretraining conditions? To resolve this inconsistency, we compared models pretrained on three different phenotype prediction tasks—age, sex, and BMI—and fine-tuned each for AD and MCI detection, examining their internal representations both before and after fine-tuning (Fig. 4A).

**Figure 4:**
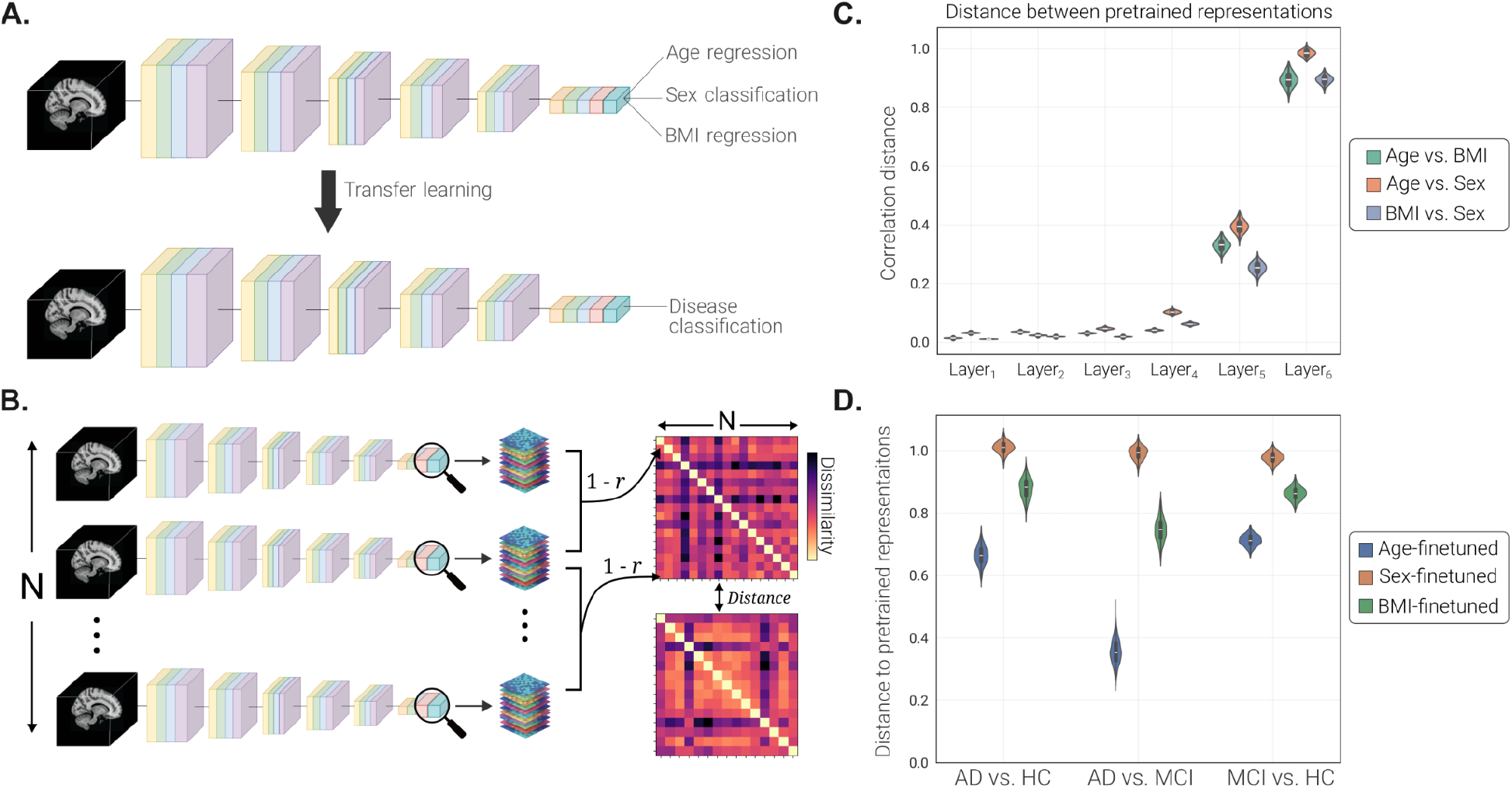
**A**. Schematics of the phenotype-estimation pretraining and fine-tuning on disease classification. **B**. RSA pipeline: given a set of brain images, Representational Dissimilarity Matrices (RDMs) are computed for each layer by calculating the dissimilarity (in this case, 1 - Pearson’s correlation coefficient) between the activations of each possible pair of brain images. These RDMs are then compared layer-wise between models using a distance measure (1 - Pearson’s correlation coefficient). **C**. Mean distance distribution of the bootstrapped RDMs distributions resulting from the test set of diagnostic tasks (matched by age and sex) in the convolutional layers of age, sex, and BMI estimation models. **D**. Distance distributions from bootstrapping a thousand times the RDMs of the last convolutional layer of fine-tuned models and their corresponding pretrained model in each diagnostic task.

The resulting models achieved state-of-the-art performance in their respective domains, with age and BMI estimation achieving a mean MAE of 2.3 (r = 0.96, p < 0.001) and 2.24 (r = 0.74, p < 0.001), respectively, and sex classification exhibiting perfect discrimination with a mean AUC-ROC of 1.0 on the test set. Subsequently, these pretrained models underwent fine-tuning using 80% of the Diseased dataset (Section 2.1) for each of the three binary classification tasks (AD vs. HC, AD vs. MCI, MCI vs. HC). A non-pretrained baseline model was also included for comparative analysis. To assess the stability of the learned representations, performance was measured using the weights from the last twenty epochs on the remaining 20% of the data (comprising 109 AD/391 HC, 113 AD/300 MCI, and 297 MCI/385 HC for the respective tasks).

After fine-tuning the phenotype-pretrained models on the three diagnostic tasks, we found that while pretraining enhanced overall robustness, the specific choice of pretraining task did not yield significant differences in diagnostic accuracy (Appx. Fig. A3). We investigated the representational basis of this convergence using representational similarity analysis^40^ (Fig. 4 B) on a subset of test images matched by age and sex (n=212, 220, and 563 for the three tasks, respectively; see Methods). We observed high similarity between the pretrained models in their first four layers (mean distance across layers 1-4: < 0.03, 0.02, 0.05, 0.10), indicating convergence towards a shared, brain-like representation regardless of the specific phenotype estimation task (Fig. 4 C). In contrast, the non-pretrained model diverged from this shared structure: its features differed significantly from those of the fine-tuned models in the first two layers (mean differences for the three tasks: 0.01, 0.09, 0.12 in layer 1; 0.07, 0.04,0.08 in layer 2; p < 0.001), and remained more similar to the raw input in the second layer (differences:-0.03, -0.04, -0.02; p < 0.001; Appx. Fig. A4 A, B).

We then explored the extent to which these brain representations shifted in the last layer after fine-tuning relative to their pretrained counterparts (Fig. 4 D). Consistent with the invariance analysis, the age prediction model exhibited the smallest deviation from its pretrained representations after fine-tuning (distance: 0.67 ± 0.03 in AD vs HC, 0.35 ± 0.03 in AD vs MCI, and 0.71 ± 0.02 in MCI vs HC), followed by the BMI prediction model (0.88 ± 0.02, 0.75 ± 0.04, 0.86 ± 0.02, respectively) and, finally, the sex classification model (1.00 ± 0.02, 0.99 ± 0.02, 0.98 ± 0.02, respectively). Notably, the largest disparity in these shifts was observed in the AD vs. MCI task, where age plays a paramount role in disentangling disease progression (see Appx. Fig. A5, A6, and A7 for additional similarity comparisons across models).

In sum, this analysis suggests that all phenotype estimation tasks drive networks to learn similar low-level features—edge detection, texture patterns, and basic anatomical structures—that generalize across tasks. The divergence in higher layers reflects task-specific specializations. During fine-tuning for disease detection, these higher layers undergo substantial reorganization, but, crucially, the age prediction model shifted the least. This pattern is consistent with the brain aging hypothesis: the age prediction model is already closer to its fine-tuned state compared to the others. The practical implication is that while pretraining on any brain-based task improves representation stability and robustness by establishing a structural scaffold, the specific choice of pretraining task is secondary to the simple benefit of having observed brain image structure.

## 3. Discussion

In this work, we explored in depth a central assumption of a predominant hypothesis in the field of neuroscience: that neurodegenerative disorders manifest as accelerated aging. This brain aging hypothesis sits at the center of current efforts in developing biomarkers from brain images for the detection of neurodegenerative diseases and yet, strikingly, it remained largely untested. Through systematic manipulation of the information learned from brain images, we causally demonstrated the inseparability of neurodegenerative phenotypes and their temporal dynamics. Our analysis indicates that aging operates as a manifold, with neurodegeneration depicting distinct patterns from healthy aging. This suggests that the necessity of aging information for neurodegeneration detection stems from the disentangling of these aging patterns and deviates from the oversimplifying conceptualization of neurodegeneration as simply accelerated aging. This intrinsic dependence on age modeling is also mirrored in the internal dynamics of neural networks, where learning disease detection drives a pronounced shift toward age-like features in later layers.

The removal of age information in our tests comes at a cost: there is a trade-off between maximizing reconstruction capacity and enforcing constraints on its representation. In the age-invariant case, we retained some age information to preserve this reconstruction capability. This was a major reason to include the age-aware model in the analyses, as it incorporates its own constraint on the representation thus closely matching the reconstruction error of the age-invariant case. It is therefore telling that, when applied to disease detection, not only did age-invariant representations perform worse than the others, but the performance of the age-aware representation was consistently similar to the age-agnostic case, despite its degraded anatomical reconstruction capability. Low-dimensional inspection of its underlying representation revealed how intertwined AD status was with aging within it: AD patients showed a shifted distribution, consistent to some extent with the accelerated aging picture, but simultaneously tighter relative to the HC and MCI populations, suggestive of a consistent representation in this space.

Our findings carry important implications for biomarker discovery in neurodegeneration. Brain age gap (BAG) has long been proposed as a suitable candidate^9,11,12^, yet recent evidence suggests subpar performance for neurodegeneration detection^27^ without a clear mechanistic explanation. By demonstrating that aging information is necessary for neurodegeneration detection, and that neurodegenerative aging deviates from—rather than accelerates—healthy aging, we provide a theoretical basis for why BAG is fundamentally ill-posed as a biomarker: it is, by construction, unidimensional and age-invariant. Moreover, these results reconcile apparently contradictory findings in the literature and refine our understanding of the brain aging hypothesis’ implications for transfer learning. Age information is causally necessary for optimal neurodegeneration detection, validating its core premise. However, this necessity doesn’t mandate explicit age-based pretraining, because aging signatures are strongly and accessibly encoded in brain morphology. In our transfer learning experiments, the single clearest observation was that representations across pretext models differ only in the top layers, with the first layers capturing structures strongly shared across tasks. This partly explains their similar performance when fine-tuned for AD detection: the first layers are task-independent. Moreover, they are also age-agnostic in the sense that age information was not removed from them, and they all remain close to the age-prediction model, so any required aging information is readily available for downstream application. A second key observation is that, across all tasks, the top convolutional layers change in a manner consistent with the importance of aging information: the age-pretrained model is modified the least of all three, and the representations of the sex-pretrained and BMI-pretrained models shift closer to the age-estimation model. These observations collectively highlight the role that aging information plays in AD models not explicitly constructed to represent it, and explain both the success of age-based transfer learning approaches^16,17^ and the finding that other pretraining strategies can achieve similar results, or confer no advantage at all^26,27^.

Our finding that age information cannot be removed from Alzheimer’s Disease (AD) detection without substantial performance loss suggests a deeper relationship between aging and neurodegeneration than previously appreciated. Unlike diseases such as Huntington’s, where age serves as a trigger but pathology follows disease-specific trajectories^41^, AD appears fundamentally inseparable from ongoing aging processes. The fact that AD divergence occurs along dimensions that also describe healthy aging variation suggests that AD may represent a pathological trajectory through multidimensional aging space rather than a discrete disease state superimposed on aging. This interpretation is consistent with mounting evidence that AD shares cellular and molecular mechanisms with healthy aging—including senescence^42^, chronic inflammation^43^, proteostasis decline^44^, and DNA damage accumulation^45^—and may help explain why anti-aging interventions show promise for neurodegeneration. In practical terms, this implies we must move beyond the constraints of unidimensional summaries like BAG that strip away the multifaceted nature of brain aging. Future work should instead prioritize multidimensional biomarkers that leverage, rather than discard, the inextricable structural coupling between age and disease phenotypes.

Several limitations of the present study should be noted. The framework models aging from a cross-sectional perspective. This is appropriate in the sense that it measures cumulative processes, but does not actually track longitudinal change over time. This is particularly relevant for our learned estimations of longitudinal change. We do not claim these estimations represent actual change in the population; they are a measure of the aging signatures that the model learned. Relatedly, in our estimation of patterns of aging, it is difficult to distinguish putative aging signals from overall anatomical heterogeneity. We attempted to mitigate these effects by decomposing not the reconstructed brain but the relative longitudinal differences, limiting the influence of absolute voxel intensities. In practice, this limitation is shared with longitudinal analyses and findings in other imaging modalities^46^. Our study also uses coarse diagnostic labels to make a general point, ignoring the heterogeneity of AD presentations and its subtypes^47^. More generally, our analyses are based on observational studies using volunteers, known to differ systematically from population studies.

Our experiments show that age information is a necessary element in AD detection (without it, performance collapses) and a sufficient one in the sense that, when present and shaping representations, models perform similarly regardless of pretraining task. The direct implication is that the AD phenotype is only defined within a coordinate system that preserves a sense of observed progression, and removing it renders the phenotype itself undefined.

## 4. Materials and Methods

### 4.1 Data Collection

We analyzed T1-weighted magnetization-prepared rapid gradient-echo (T1-MPRAGE; 1mm^3^) brain MRI scans collected from several sources, following a streamlined workflow optimized for efficiency and large-scale processing^48^ (details in Section 4.1.1). We retained scans from participants older than 20 and younger than 95. The remaining images were split in two: the *general population* dataset, composed of brain images collected from the general population, and the *diseased* dataset, composed of brain images that gather Alzheimer’s Disease (AD) and Mild Cognitive Impairment (MCI) patients, and their corresponding Healthy Control (HC) subjects.

#### 4.1.1 Image Preprocessing

The protocol followed^48^, using ANTsPy^50^ for image processing, which included skull stripping and intensity inhomogeneity correction. Following best practices for deep learning applications, we aligned the scans to a standardized MNI reference frame using affine transformations. Additionally, we generated non-linear deformation maps between the affine-registered images and MNI space, which facilitated the pooling and visualization of aging decomposition results.

### 4.2 Variational Autoencoders

As the input data were T1-w brain images cropped to the center (of dimensions 160 x 192 x 160), we employed 3D convolutional neural networks. In particular, we made use of the Simple Fully Convolutional Network (SFCN)^51^ for the encoder, with the exception that pooling layers were removed and the convolutional layers used stride two instead. The decoder was a mirror of the encoder that used transposed convolutions and no activation function in its final layer.

#### 4.2.1 Age-agnostic, Age-aware and Age-invariant Variations

We designed three different architectures of VAEs: agnostic to age, maximally informed about age, and maximally uninformed about age. To create brain representations maximally informed about age, age was estimated from the encodings through a linear regression during training of the VAE. To remove age information, we made use of the Invariant Conditional VAE (ICVAE)^28^, in which the mutual information between the encoded data (z) and a sensitive attribute (age) is minimized through a variational upper bound:

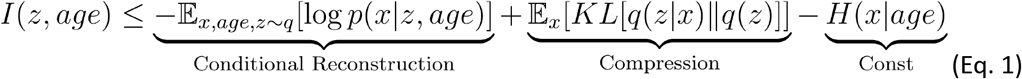

Where *x* is the input data, *p* is the decoder and *q* the encoder. This is similar to the encoder-decoder pair in VAEs, with the exception that the decoder is conditional and there is an additional KL term. The derivation for this formula can be found in^28^. Following their work, the decoder was conditioned by concatenating age as a floating-point number to the encoding *z* (see Appendix A4 for other conditioning variations).

All of these variations shared two different objective functions from the original VAE definition: the “distance” to the prior loss (under the term *β* * [*q*(*z*|*x*)| *𝒩*(0,*I*)]) and the reconstruction loss (defined as 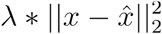, where *x* is the input and 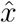 the output from the decoder). As noted in Eq. 1, the *age-invariant* model had an additional loss term (*γ* * *KL*[*q*(*z*|*x*) |*q*(*z*)]) that was approximated using a pairwise Gaussian KL divergence. , *β, λ* and *γ* are hyperparameters tuned on the validation set.

Finally, as the *age-aware* model also estimated age with a linear regression from the encoding *z* during training, a KL divergence loss term was added. Age was encoded as a Gaussian distribution with mean equal to the true value and a standard deviation of one, encompassed in a vector of size 70 (ages 20-90)^51^. The final age estimation was computed as the weighted average of these bins, where weights were derived from the estimated probability for each bin.

#### 4.2.2 Model Training

When splitting data from the General population dataset into training and validation, the difference in size between cohorts had to be taken into account (see Table 1). 10% (4,216 samples) and 15% (6,627 samples) of the UK-Biobank were split into validation and testing, respectively, whereas the rest of the datasets were split into 20% (525 samples) and 10% (214 samples) for validation and testing, respectively. In total, 33,110 different brain images were employed for model training, during which 2,000 samples (with replacement) were drawn from each of the smaller datasets to ensure a higher representation from them.

VAEs training ran for 100 epochs with a batch size of 16, using two V100 GPUs and PyTorch Lightning’s Distributed Data Parallel algorithm. AdamW was employed as the parameters’ optimizer, with an initial learning rate of 4e-4 linearly decaying to 4e-5. Hyperparameters were set by maximizing performance (reconstruction error and age estimation or invariance) in the validation set: , and (described in Section 4.2.1) were set to 1e-4, 1 and 0 for the *age-agnostic* and *age-aware* models, and to 1e-6, 3 and 1e-3, respectively, for the *age-invariant* model. The weight of the KL divergence term for age estimation in the *age-aware* model was set to 100. All three models used a latent space dimension of 250.

#### 4.2.3 Evaluation

To evaluate the information contained in the brain encoding models described in 4.2 after being trained on the General population train set, we encoded the brain images from the General population test set and used them as input to a shallow neural network and estimated their age. The validation and test sets of the General population dataset were split in 4,483 and 6,277 samples to train the feedforward network (75% from UKBB, the remainder equally resampled from the other five cohorts), and in 633 and 1,041 samples for evaluation. The validation set was further split 70:30 for hyperparameter selection of the neural network, that comprised two hidden layers (64 and 32 units), trained with AdamW (default parameters), a linearly decaying learning rate (1e-3 to 1e-4), batch size 32, and 50 epochs. Age was represented with soft labels (see Section 4.2.1) and KL divergence was used as the loss function. Mean Absolute Error (MAE) and Pearson’s correlation coefficient between the predicted and true values were used for measurement. To measure the reconstruction error, we computed the Mean Squared Error (MSE) between the original and reconstructed brain images.

Discrimination between conditions (AD, MCI, or HC) followed the same methodology: brain images from the Diseased dataset were encoded using the encoders of the previously trained VAEs and these encodings were used as input to a feedforward network with two hidden layers that was trained to distinguish between either AD and HC, AD and MCI, or MCI and HC. The Diseased dataset matched by age and sex (see Section 2.2.1) was split 80:20 into training and test sets, with the training set further divided 80:20 for hyperparameter selection of the classifier. As these tasks pertain to binary classification, Binary Cross Entropy (BCE) was employed as the loss function. For the computation of the brain-age-gap (BAG) of individuals, the difference between chronological and estimated brain age was computed by the model from^51^ trained on the General Population dataset.

In both cases, a bootstrapping strategy was employed on the testing set of the classifier, conducting random sampling with replacement a thousand times, and statistical significance between distributions was determined by the proportion of lower values (given that bootstrapping samples are not independent from each other and statistical tests, such as Wilcoxon’s rank-sum test, would be ill-posed).

### 4.3 Simulation and Decomposition of Aging Patterns

To analyze aging processes, we selected a random subsample of participants from the General population test set, comprising 300 young (aged 45–60) and 200 old (aged 70–85) individuals. We reconstructed expected scans for each participant at extreme ages using the conditional decoder described in Section 4.2.1: 85 years for the young group and 45 years for the old group. For each individual, we computed a relative difference map between the expected and actual reconstructions. This was calculated as the voxelwise ratio of the difference to the sum voxel intensities, yielding a value bounded between -1 and 1 to capture both the direction and magnitude of age-related change. To perform linear decomposition, these difference maps were first normalized and transformed to a common MNI152 space.

To prepare the data for NMF, we explicitly isolated the negative values from the difference maps (representing tissue loss) and converted them to absolute values. Following the decomposition into three components, we applied L2 normalization and thresholded the maps to remove arbitrary low values (<10^-4^). Voxel-wise assignment for visualization was performed based on the component with the highest coefficient value.

To compare aging patterns in pathology, we selected a cohort of elderly participants (over 75 years old), consisting of 300 patients with Alzheimer’s Disease (AD) and 280 healthy controls (HC), matched for age and sex. For each individual, we generated a reconstruction targeted at age 45. We computed the difference maps following the procedure described above and projected them onto the established NMF components. Statistical differences in component weight distributions between the AD and HC groups were assessed using a Wilcoxon rank-sum test.

### 4.4 Model Pretraining and Fine-tuning

Phenotype (age, sex, or BMI) estimation was carried out using SFCN^51^ as the model backbone and followed the same hyperparameters of that study. Age estimation was carried out in the same way as described in Section 4.2.1 for the age-aware model. Likewise, BMI estimation was performed on the 10-60 range with a vector of size 50. Given that BMI labels were not present in all cohorts of the General population dataset, only those brain images from UKBB, HCP and HCP-aging were employed (totaling 32,192 different brain images). Model hyperparameters followed those of^51^ and training ran for 130 epochs.

Disease detection fine-tuning employed the same hyperparameters as those for pretraining and ran for a hundred epochs. Since the number of parameters to train was considerably bigger than those of the shallow neural network in Section 4.2.3, inverse probability weights (IPW) in conjunction with a cut-off based on a Gaussian fit on the distribution (higher than 0.4, resulting in a 56-84 range) were used to control for age confounding instead of constraining the size of the dataset.

#### 4.4.1 Representational Comparison between Pretrained and Fine-tuned models

Representational Similarity Analysis (RSA) was carried out primarily in two steps. First, a forward pass was done in each pretrained and fine-tuned model, where the features (activations) from each layer for each brain image were extracted. A within-layer comparison of each pair of instances was then performed by computing the dissimilarity between their activations (one minus Pearson’s correlation coefficient). For a set of *N* brain images, this yielded an *NxN* representational dissimilarity matrix (RDM) that depicted how (dis)similar the layer activations were for each pair of brain images. The second step consisted of a layer-wise comparison between models by computing Pearson’s correlation coefficient between their corresponding RDMs: the higher it was, the more similar their representations. This second step was bootstrapped a thousand times to obtain a similarity distribution. For the comparison across modalities (pretrained and non-pretrained), bootstrap distributions were averaged within each modality and subtracted; significance was estimated as the proportion of values above zero after mean-centering the difference distribution.

## Data Availability

The data used in this study are publicly available from the respective data owners.

## Code Availability

The source code for model training, testing and analysis can be found in https://github.com/NeuroLIAA/brain-icvae.

## Acknowledgments

The datasets analyzed in this study were sourced from the Alzheimer’s Disease Neuroimaging Initiative (ADNI) database (adni.loni.usc.edu); the Australian Imaging Biomarkers and Lifestyle Study of Ageing (AIBL, www.aibl.csiro.au); the Cambridge Centre for Ageing and Neuroscience (CamCAN) repository (www.mrc-cbu.cam.ac.uk/datasets/camcan/); the HCP Young Adult and HPC Aging studies (ida.loni.usc.edu/login.jsp); OASIS (www.oasis-brains.org); and the UK Biobank Resource (www.ukbiobank.ac.uk/), under Application Number 95318.

The HCP project (Principal Investigators: Bruce Rosen, M.D., Ph.D., Martinos Center at Massachusetts General Hospital; Arthur W. Toga, Ph.D., University of Southern California, Van J. Weeden, MD, Martinos Center at Massachusetts General Hospital) is supported by the National Institute of Dental and Craniofacial Research (NIDCR), the National Institute of Mental Health (NIMH), and the National Institute of Neurological Disorders and Stroke (NINDS). HCP is the result of efforts of coinvestigators from the University of Southern California, Martinos Center for Biomedical Imaging at Massachusetts General Hospital (MGH), Washington University, and the University of Minnesota. HCP data are disseminated by the Laboratory of Neuro Imaging at the University of Southern California.

Data collection and sharing for the ADNI project was funded by ADNI (National Institutes of Health Grant U01 AG024904) and DOD ADNI (Department of Defense award number W81XWH12-2-0012). ADNI is funded by the National Institute on Aging, the National Institute of Biomedical Imaging and Bioengineering, and through generous contributions from the following: AbbVie, Alzheimer’s Association; Alzheimer’s Drug Discovery Foundation; Araclon Biotech; BioClinica, Inc.; Biogen; BristolMyers Squibb Company; CereSpir, Inc.; Cogstate; Eisai Inc.; Elan Pharmaceuticals, Inc.; Eli Lilly and Company; EuroImmun; F. Hoffmann-La Roche Ltd and its affiliated company Genentech, Inc.; Fujirebio; GE Healthcare; IXICO Ltd.; Janssen Alzheimer Immunotherapy Research & Development, LLC.; Johnson & Johnson Pharmaceutical Research & Development LLC.; Lumosity; Lundbeck; Merck & Co., Inc.; Meso Scale Diagnostics, LLC.; NeuroRx Research; Neurotrack Technologies; Novartis Pharmaceuticals Corporation; Pfizer Inc.; Piramal Imaging; Servier; Takeda Pharmaceutical Company; and Transition Therapeutics. The Canadian Institutes of Health Research is providing funds to support ADNI clinical sites in Canada. Private sector contributions are facilitated by the Foundation for the National Institutes of Health (www.fnih.org). The grantee organization is the Northern California Institute for Research and Education, and the study is coordinated by the Alzheimer’s Therapeutic Research Institute at the University of Southern California. ADNI data are disseminated by the Laboratory for Neuro Imaging at the University of Southern California.

The authors express their gratitude to Juan E. Kamienkowski and Ricardo F. Allegri for revising the manuscript; Matías Aiskovich for their work in data preparation and processing; Gustavo Stolovitsky for encouraging this work; and Kenny Ng for support with UK Biobank.

## Ethics statement

The current study involves the secondary analysis of de-identified, publicly available datasets. The original data collection for all contributing cohorts (ADNI, AIBL, Cam-CAN, HCP Young Adult, HCP Aging, OASIS, and UK Biobank) was conducted in accordance with the Declaration of Helsinki and was approved by the respective institutional review boards (IRBs) or ethics committees at each participating site. Informed consent was obtained from all participants or their legal guardians at the time of original enrollment.

## Author contributions

**FT:** Methodology, Software, Visualization, and Writing – original draft. **AM:** Software and Writing – review & editing. **EC:** Validation, and Writing – review & editing. **HL:** Software and Writing – review & editing. **JR:** Supervision and Writing – review & editing. **AD:** Supervision and Writing – review & editing. **PMR:** Writing – review & editing. **DFS:** Writing – review & editing. **GAC:** Conceptualization, Supervision, and Writing – review & editing. **PP:** Conceptualization, Supervision, and Writing – original draft.

