## Appendix for "Alzheimer’s Disease Brain Phenotypes are Age-dependent"

### A1. Age distribution in Diseased dataset

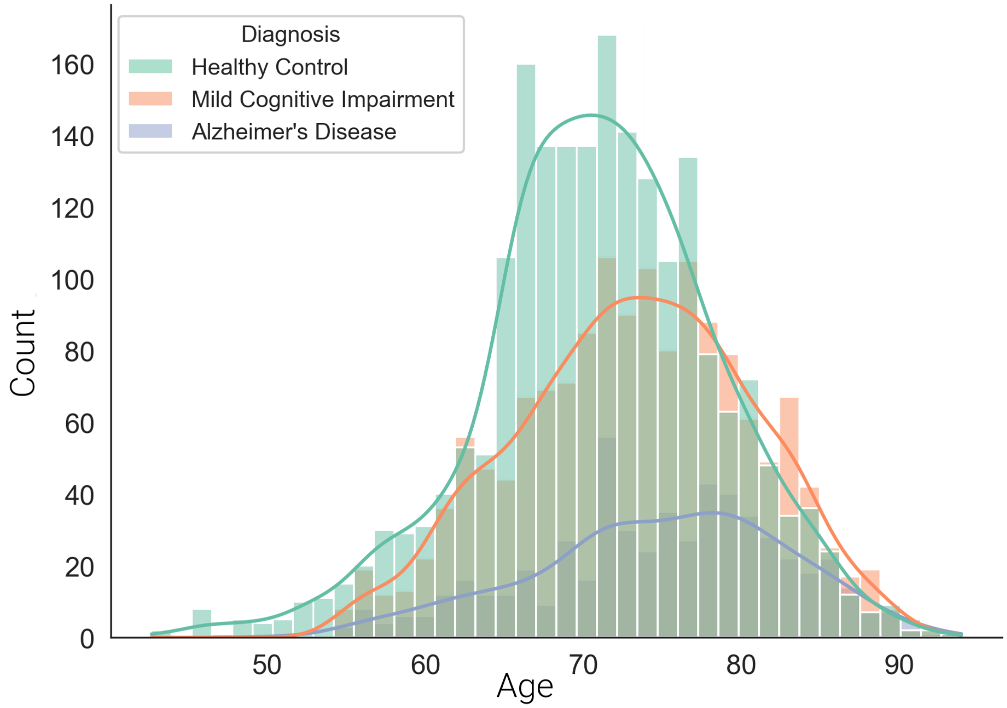


*Figure A1: Age distribution colored by diagnosis in the Diseased dataset.*

### A2. Controlling for other phenotypic information

To guarantee our results are about age information and not due to some other unobserved variable, we introduce three control models: a *sex-invariant* model, a *sex-aware* model and a *BMI-aware* model (given that BMI information was not available in all cases, no *BMI-invariant* model was developed). The *sex-invariant* model follows the same architecture as the *age-invariant* model, except that the decoder is conditioned on sex (a binary label). In the case of *sex-* and *BMI-aware*, they are analogous to the *age-aware* model but estimating sex and BMI, respectively, from the encodings instead of age during training. As the variable *sex* is represented in the dataset as a binary label, Binary Cross Entropy (BCE) was used as the loss function when predicting it in the *sex-aware* model and the baseline reported corresponds to random labels from a binomial distribution whose probability equals the proportion of the majority class label. BMI, on the other hand, is represented with soft labels in the same way as age with a vector of size 46 (BMI 13-59) (see Section 4.1.2.) and the Kullback-Leibler divergence is used as the loss function. Additionally, when the data were missing no prediction was made.

Below we report the performance of the resulting brain representations in predicting age, sex and BMI as well as their reconstruction and disease classification performance as described in Section 2.2, along with the aging variants.

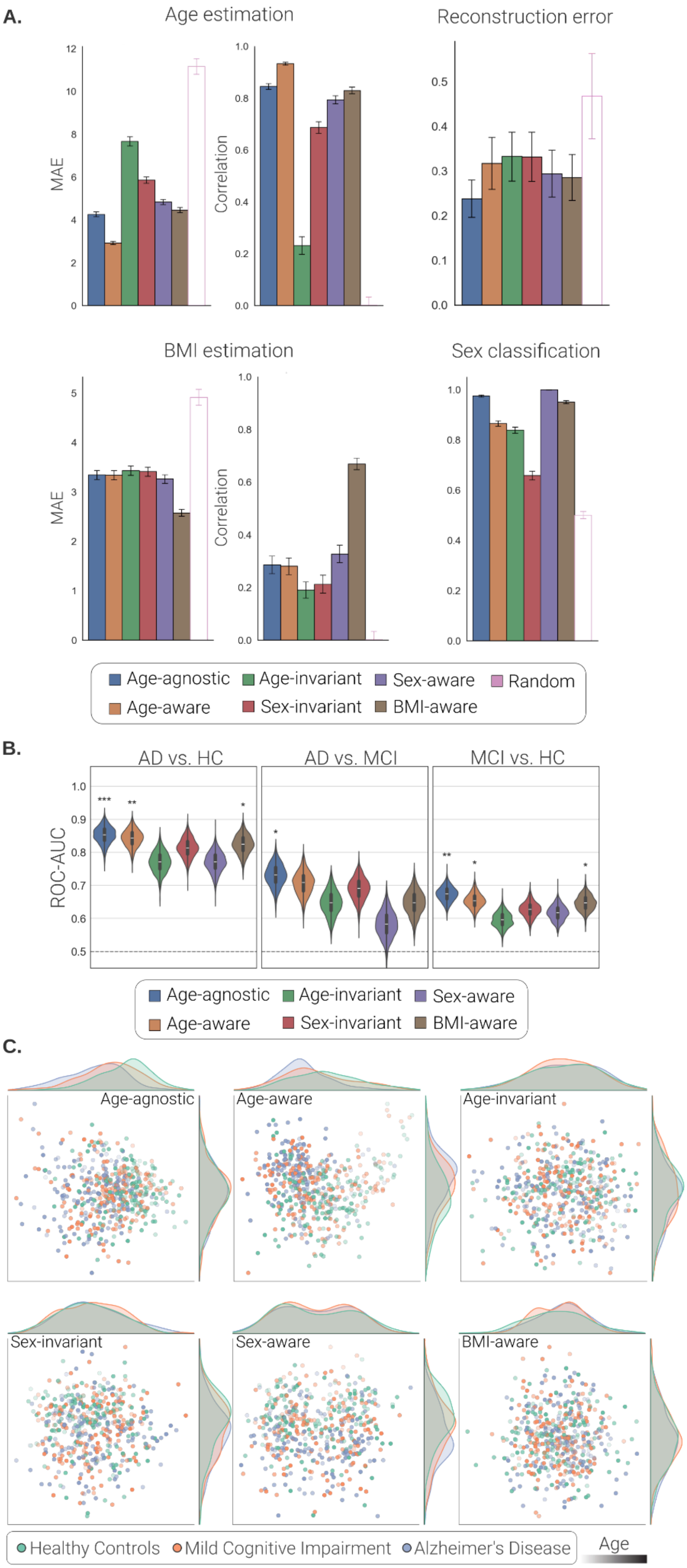
A.
Age estimation
Reconstruction error
MAE
Correlation
BMI estimation
Sex classification
Age-agnostic
Age-invariant
Sex-aware
Random
Age-aware
Sex-invariant
BMI-aware
B.
AD vs. HC
AD vs. MCI
MCI vs. HC
ROC-AUC
C.
Age-agnostic
Age-aware
Age-invariant
Sex-invariant
Sex-aware
BMI-aware
Healthy Controls
Mild Cognitive Impairment
Alzheimer's Disease
Age

Figure A2: **A.** Phenotype estimation and reconstruction performance from the different brain representations (age-invariant, age-agnostic, age-aware, sex-invariant, sex-aware and BMI-aware) on the General population dataset (2.1). Error in numeric variables, such as age and BMI, is quantified using Mean Absolute Error (MAE) and Pearson's correlation. The random baseline is constructed by shuffling the labels. Sex prediction is measured by the AUC of the ROC curve, and the random baseline consists of a random sampling from a Bernoulli distribution with *p* equal to the proportion of the majority class label. The reconstruction error corresponds to the Mean Squared Error (MSE) between the input (T1-w images) and the output of the decoder (i.e., the reconstructed T1-w images). **B.** Performance on the detection of Alzheimer's Disease against Healthy Controls (first from the left), Alzheimer's Disease and Mild Cognitive Impairment (middle), and Mild Cognitive Impairment and Healthy Controls (third from the left) using the different variations of brain representations, bootstrapped a thousand times. The violin plots reflect the Kernel Density Estimate (KDE) and the median and interquartile range of the ROC-AUCs distribution. Asterisks above the violins indicate statistical significance against age-invariant, determined by the proportion of lower values. **C.** First two principal components on a random 40% (N=678) sample of the matched brain images. Marker size is defined by the age of the individual.

As expected, the "aware" variations achieve the greatest performance in their respective domains (Fig. A2 A and Table A1). BMI-aware achieves a 2.58 mean MAE (std = 0.07; r = 0.67, $p < 0.001$) in BMI estimation, Sex-aware achieves a mean ROC-AUC of 1.0 (std = 0.0007) in sex classification, and Age-aware achieves a 2.92 mean MAE (std = 0.07; r = 0.93, $p < 0.001$) in age estimation. However, BMI-aware and Sex-aware have comparable reconstruction errors (0.29 mean MAE, std = 0.05) that are significantly better ($p < 0.01$) than those achieved by Age-aware (0.32 mean MAE, std = 0.06), potentially due to the lower values of their respective loss functions. Conversely, Sex-invariant yields the lowest performance in sex classification (0.66 mean ROC-AUC, std = 0.02), with comparable reconstruction performance to Age-invariant and Age-aware (0.33 mean MSE, std = 0.06), but displaying significantly better ($p < 0.001$) age estimation than the former (5.86 mean MAE, std = 0.15; r = 0.69, $p < 0.001$).

Table A1: Age, BMI, and sex estimation, and the reconstruction error on the test set of the General population dataset corresponding to the different variations of brain representations.

| | **Age** | | **BMI** | | **Sex** | **Recons. Error** |
|---|---|---|---|---|---|---|
| | **MAE** | **Corr.** | **MAE** | **Corr.** | **ROC-AUC** | **MSE** |
| **Age-agnostic** | 4.26 (±0.11) | 0.84 | 3.34 (±0.09) | 0.28 | 0.97 (±0.00) | 0.24 (±0.04) |
| **Age-aware** | 2.92 (±0.07) | 0.93 | 3.34 (±0.09) | 0.28 | 0.86 (±0.01) | 0.32 (±0.06) |
| **Age-invariant** | 7.66 (±0.21) | 0.23 | 3.43 (±0.09) | 0.19 | 0.84 (±0.01) | 0.33 (±0.05) |
| **Sex-invariant** | 5.86 (±0.15) | 0.69 | 3.41 (±0.09) | 0.21 | 0.66 (±0.02) | 0.33 (±0.06) |
| **Sex-aware** | 4.83 (±0.13) | 0.79 | 3.26 (±0.09) | 0.33 | 0.99 (±0.00) | 0.29 (±0.05) |
| **BMI-aware** | 4.45 (±0.12) | 0.83 | 2.58 (±0.07) | 0.67 | 0.95 (±0.01) | 0.28 (±0.05) |
| **Random** | 11.16 (±0.37) | 0.00 | 4.92 (±0.16) | 0.00 | 0.5 (±0.02) | 0.47 (±0.09) |

When observing the differences in disease classification performance of the different variations of brain representations (Fig. A2 B and Table A2), some clear patterns emerge. On one hand, sex information appears to confound disease detection, as the *sex-aware* brain representations perform poorly in all three classification tasks, whereas its counterpart (*sex-invariant)* actually performs better in spite of having a higher reconstruction error and less age information (Table A1). *BMI-aware* brain representations, on the other hand, in spite of possessing a significantly lower reconstruction error than *age*-aware, fails to outperform it, indicating the relevance of the addition of age information for these tasks.

Table A2: Mean ROC-AUC and interquartile range of the different variations of brain representations in the disease classification tasks.

| | AD vs. HC | AD vs. MCI | MCI vs. HC |
|---|---|---|---|
| **Age-agnostic** | 0.85 (0.03) | 0.73 (0.05) | 0.67 (0.03) |
| **Age-aware** | 0.84 (0.03) | 0.71 (0.05) | 0.65 (0.03) |
| **Age-invariant** | 0.77 (0.04) | 0.65 (0.05) | 0.59 (0.03) |
| **Sex-invariant** | 0.81 (0.04) | 0.69 (0.05) | 0.63 (0.03) |
| **Sex-aware** | 0.77 (0.04) | 0.58 (0.05) | 0.62 (0.03) |
| **BMI-aware** | 0.82 (0.04) | 0.65 (0.05) | 0.65 (0.03) |

### A3. Phenotype estimation pretraining and fine-tuning

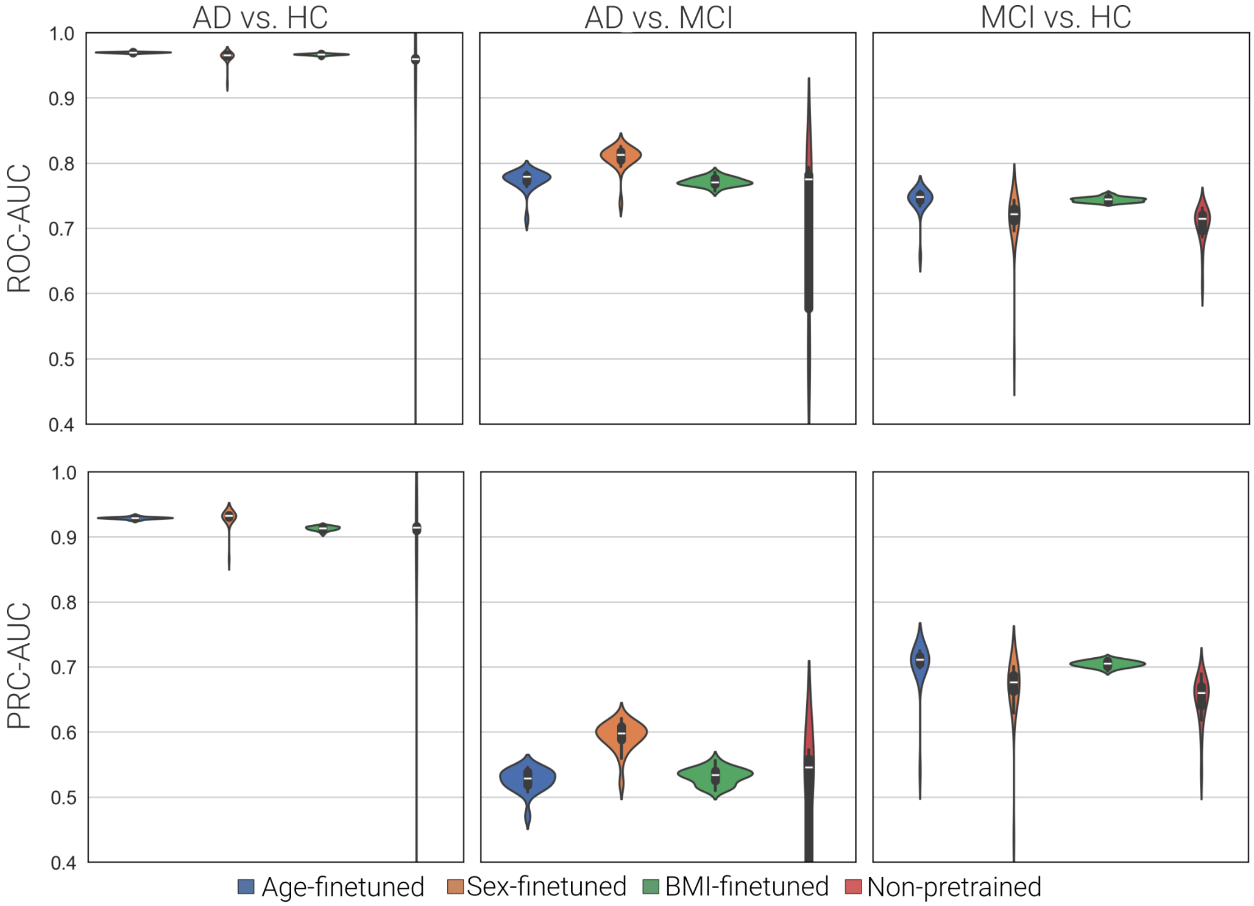


**Figure A3:** Performance on the detection of Alzheimer's Disease against Healthy Controls (first from the left), Alzheimer's Disease and Mild Cognitive Impairment (middle), and Healthy Controls and Mild Cognitive Impairment (third from the left) of the phenotype-pretrained and non-pretrained models as measured by ROC- (top) and PRC-AUCs (bottom) resulting from the weights of the last twenty epochs. The violin plots reflect the Kernel Density Estimate (KDE) and the median and interquartile range of the distributions.

Figure A3 depicts the ROC (top) and precision-recall (bottom) curves AUCs of the fine-tuned pretrained and non-pretrained models on the disease classification tasks. Both the training and testing sets belong to the same composite dataset, so these results are not externally validated. After fine-tuning, no significant differences were observed between the pretrained models across the pathology discrimination tasks. The age-pretrained model achieved mean ROC-AUC and PRC-AUC values of 0.97

and 0.93 (IQR: 0.001) for AD vs. HC, 0.77 and 0.53 (IQR: 0.009) for AD vs. MCI, and 0.70 and 0.70 (IQR: 0.008) for MCI vs. HC. The sex- and BMI-pretrained models showed comparable performance across these tasks (Sex: 0.96/0.93, 0.81/0.59, 0.74/0.66; BMI: 0.97/0.91, 0.77/0.53, 0.74/0.70). Conversely, the non-pretrained baseline yielded similar averages (0.93/0.88, 0.81/0.47, 0.74/0.65), yet its performance was notably less stable and exhibited more outliers.

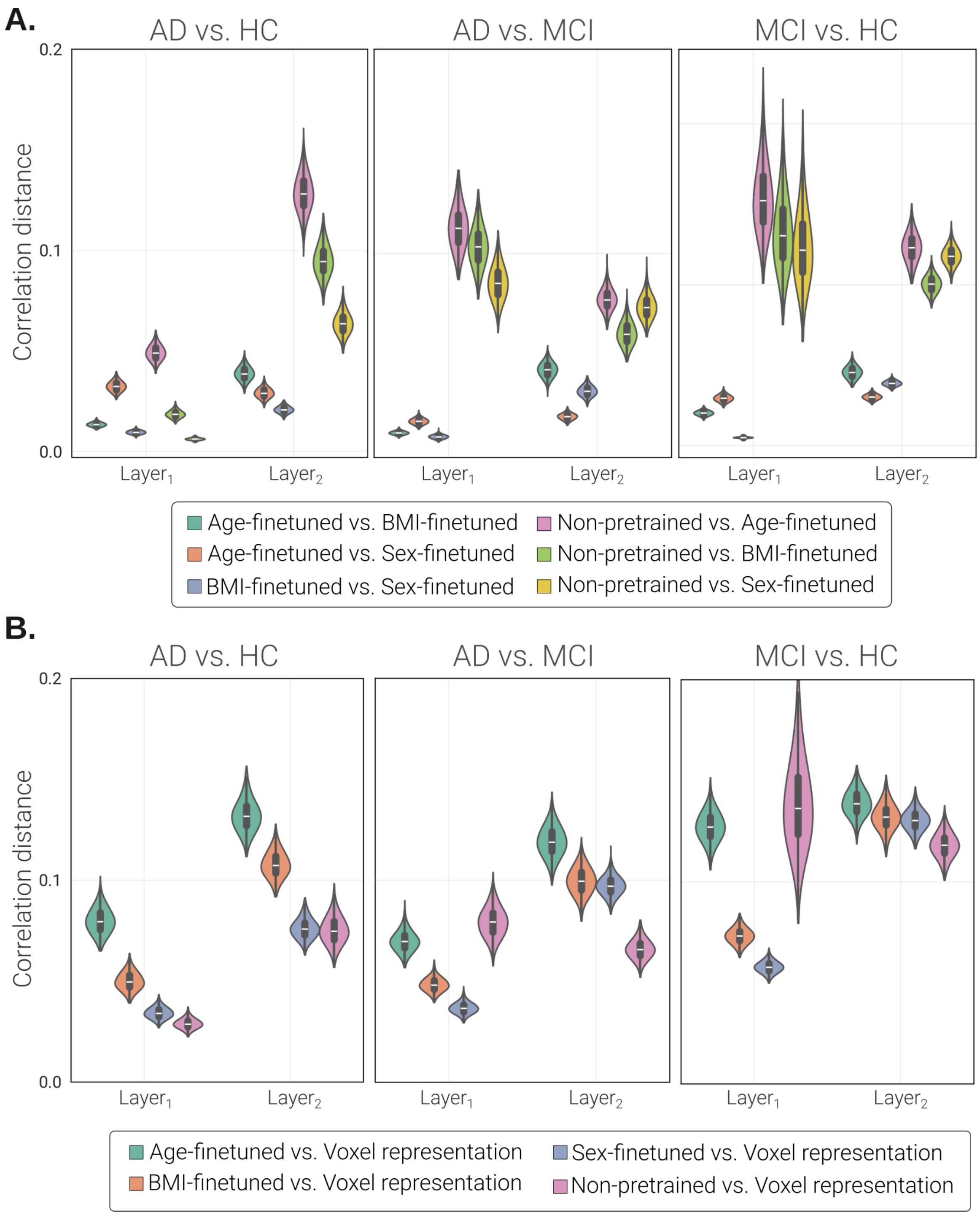

Figure A4: **A.** Representational dissimilarity distributions on the first two layers of each of the fine-tuned models and the non-pretrained model on the disease classification tasks matched by age and sex (N=212, 220 and 563 for AD versus HC, AD versus MCI, and MCI versus HC, respectively). **B.** Representational dissimilarity distributions on the first two layers of each of the fine-tuned models and the non-pretrained model against RDMs built on the raw input (*Voxel representation*) on the disease classification tasks matched by age and sex (N=212, 220 and 563 for AD versus HC, AD versus MCI, and MCI versus HC, respectively).

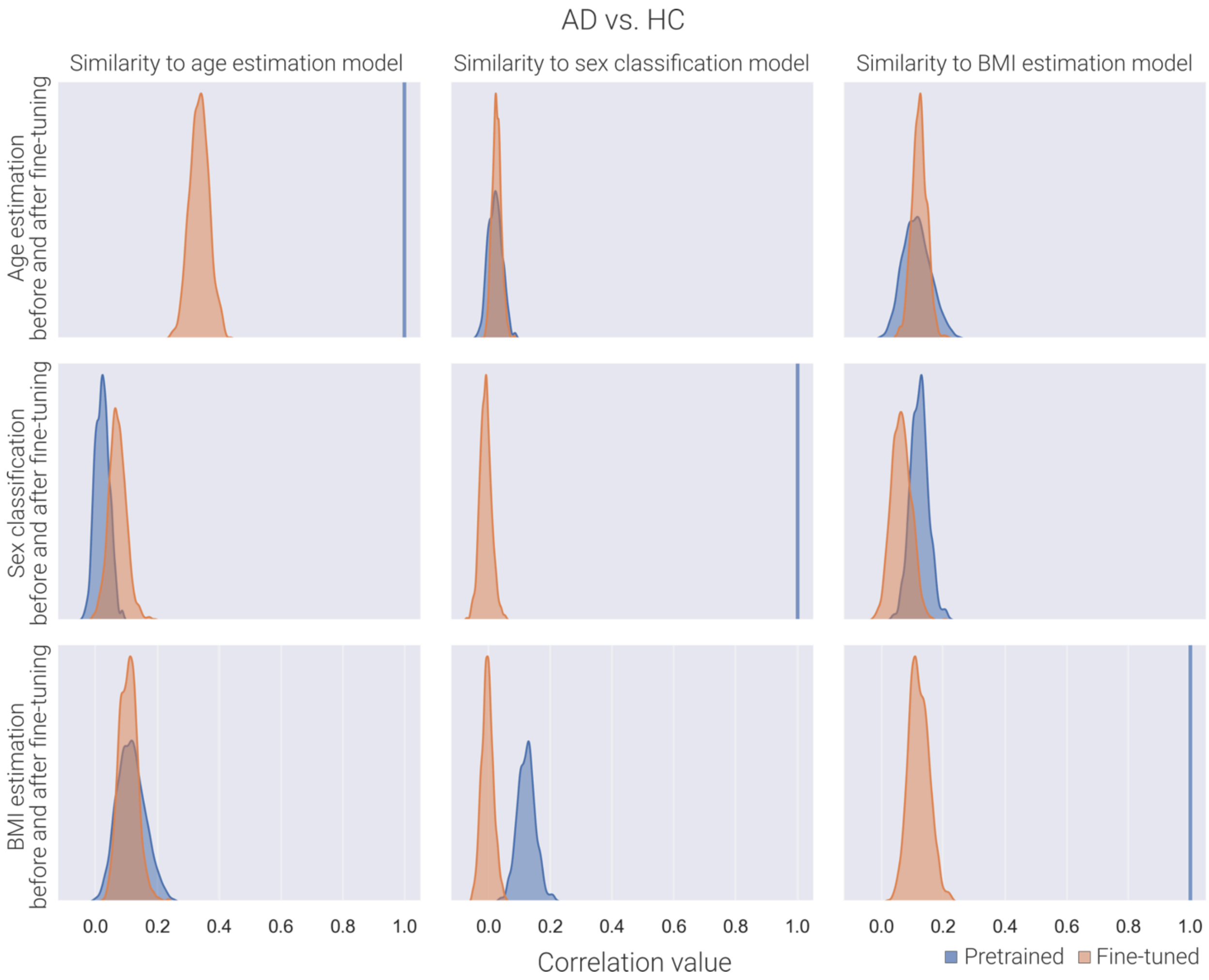


Figure A5: Similarity densities between the representations in the last convolutional layer of pretrained and fine-tuned models on the AD versus HC task, resulting from bootstrapping the RDMs a thousand times. Columns indicate different pretrained modalities (either age, sex, or BMI estimation), while rows depict how the representations of each pretrained and fine-tuned model compared to that specific pretrained modality. The distance between blue and orange densities corresponds to the degree of change with respect to the representations of that specific phenotype estimation model.

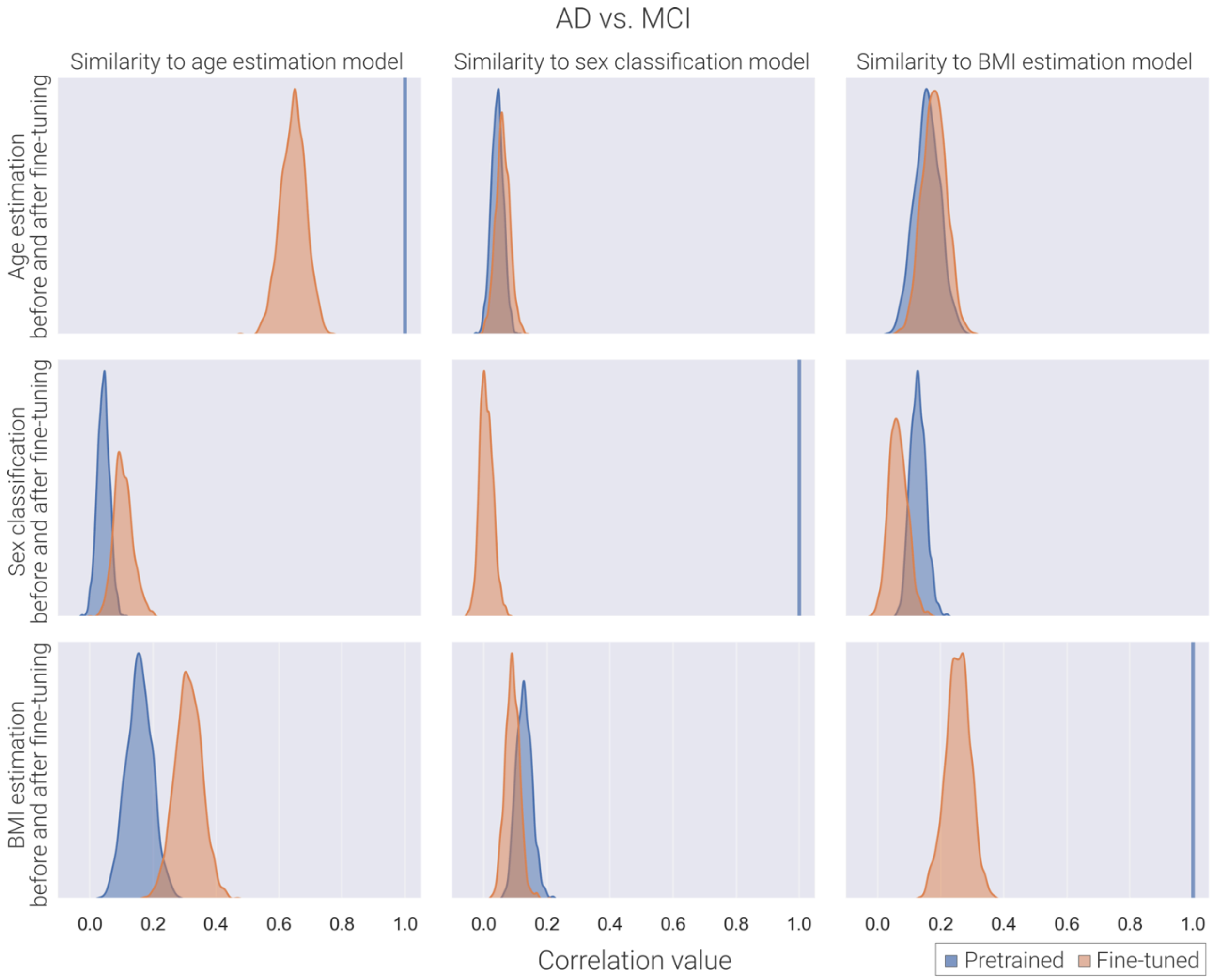


Figure A6: Similarity densities between the representations in the last convolutional layer of pretrained and fine-tuned models on the AD versus MCI task, resulting from bootstrapping the RDMs a thousand times. Columns indicate different pretrained modalities (either age, sex, or BMI estimation), while rows depict how the representations of each pretrained and fine-tuned model compared to that specific pretrained modality. The distance between blue and orange densities corresponds to the degree of change with respect to the representations of that specific phenotype estimation model.

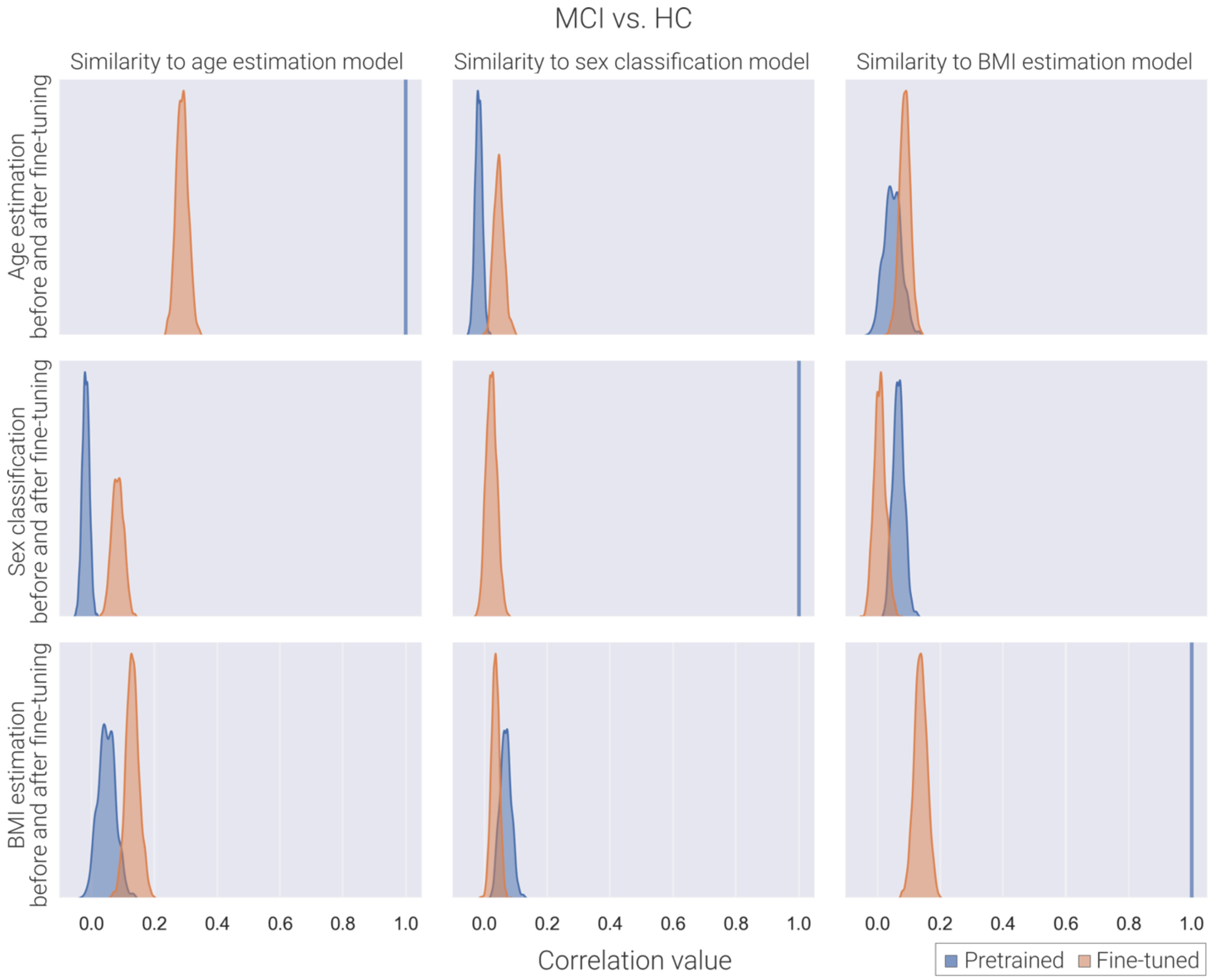


Figure A7: Similarity densities between the representations in the last convolutional layer of pretrained and fine-tuned models on the MCI versus HC task, resulting from bootstrapping the RDMs a thousand times. Columns indicate different pretrained modalities (either age, sex, or BMI estimation), while rows depict how the representations of each pretrained and fine-tuned model compared to that specific pretrained modality. The distance between blue and orange densities corresponds to the degree of change with respect to the representations of that specific phenotype estimation model.

### A4. Age conditioning variations

Conditioning of the conditional decoder presented in Section 2.2 can be achieved with different representations of the numerical value to condition with. In the case of age, we explored three possible variations: as a floating-point number, as a soft label (a vector of size 70, see Section 4.2.1) or as a sinusoidal vector (equivalent to the positional encodings used in transformers). In this last case, the value is not concatenated to the latent vector, but rather it is summed.

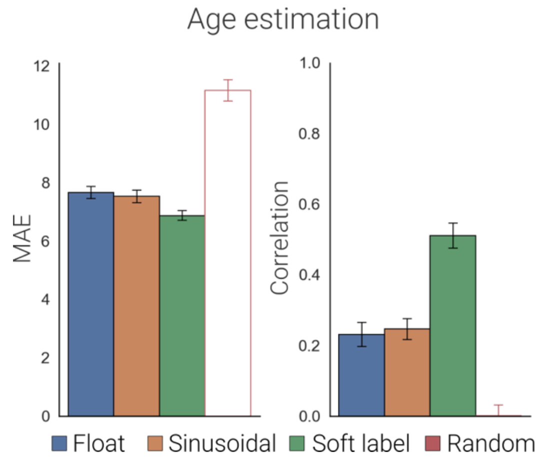


Figure A8: Age estimation using the resulting brain representations from the different age conditioning variations. *Float* refers to conditioning representing age as a floating-point number, while *Sinusoidal* refers to conditioning representing age with a sinusoidal vector, and *Soft label* refers to conditioning with a vector that represents age as a Gaussian distribution.

As depicted in Figure A8, the highest removal of age information is achieved when the decoder is conditioned with a floating-point number for representing age (MAE 7.66, std. 0.21, $r = 0.23$), followed closely by the sinusoidal conditioning (MAE 7.53, std. 0.22, $r = 0.25$). Conditioning with a soft label performs the worst (MAE 6.87, std. 0.17, $r = 0.51$), potentially due to the concatenation of a large vector to the latent representations (as it represents over 30% of its final size).
